# Heterozygous Loss of Scn1b Results in Concealed Conduction Phenotype Unmasked by Osmotic Stress

**DOI:** 10.64898/2026.06.15.732501

**Authors:** Rowan Maisonneuve, Chandra Bain, Clare Dennison, Mark Warren, Gregory S Hoeker, Robert G. Gourdie, Steven Poelzing

**Author notes:** **Address for Correspondence:** Steven Poelzing, Ph.D., 2 Riverside Circle, Rm R2115, Roanoke, VA 24016. **Author Information:** Rowan Maisonneuve, Chandra Bain, Clare Dennison, Mark Warren, Gregory S Hoeker, Robert G. Gourdie.

## Abstract

**Rationale:** *SCN1B* encodes the β-subunits of the main cardiac voltage-gated sodium channel, Na_V_1.5. Variants are linked to cardiac conduction disease, often with concealed phenotypes. Whether β1-subunits regulate conduction through nanoscale intercalated disc (ID) structures, e.g. perinexi, and ephaptic coupling remains unresolved.

**Objective:** Test whether Scn1b haploinsufficiency induces latent conduction abnormalities that are unmasked by perturbations in extracellular nanodomains.

**Methods and Results:** Adult Scn1b+/− mice and wild-type (WT) littermates underwent multiscale phenotyping (qRT-PCR, Western blot, patch clamp, transmission electron microscopy (TEM), *ex vivo* optical mapping, *in vivo* ECG). Scn1b+/− hearts showed ∼50% reductions in Scn1b mRNA and β1 protein without changes in canonical conduction proteins. Peak sodium current, baseline conduction velocity *ex vivo*, and baseline QRS duration *in vivo* were unchanged. However, TEM revealed increased baseline perinexal width in Scn1b+/− hearts. Osmotic expansion of the perinexus with mannitol slowed conduction to a greater extent in Scn1b+/− hearts and prolonged QRS duration *in vivo*. In contrast, perinexal narrowing with dextran 2MDa selectively increased conduction velocity in Scn1b+/− hearts.

**Conclusions:** Scn1b haploinsufficiency preserves baseline excitability and conduction but structurally remodels the ID at the nanoscale, increasing sensitivity to extracellular nanodomain perturbations. These data support a structural role for β1-subunits in ephaptic coupling, and that conduction is maintained over a range of perinexal widths with pathological conduction slowing occurring beyond a critical width. Importantly, osmotic stress unmasks a concealed conduction phenotype, identifying extracellular nanodomain stability as a potential therapeutic target to mitigate arrhythmia risk in *SCN1B*-associated disease.

## INTRODUCTION

Sudden cardiac death is a leading cause of death globally, often due to ventricular arrhythmias.^1^ Reentrant ventricular arrhythmias are particularly fatal^2^ and arise from reduced excitability and slowed conduction, both of which depend on sodium channel function to initiate the action potential.^3^ Propagation of the action potential through the myocardium subsequently relies on either gap junction^4^ or ephaptic^5^ coupling. A proposed mediator of ephaptic coupling in cardiac tissue is the perinexus, an intercellular nanodomain located in the intercalated disc (ID).^6^ In brief, cardiac ephaptic coupling refers to the mechanism in which activation of a primary cardiomyocyte causes sodium influx from extracellular nanodomains (approximately 20nm wide) to locally deplete extracellular ions within the perinexus, generating a transient negative extracellular potential that facilitates trans-activation of sodium channels in the adjacent cell and enables action potential propagation.

Alterations in sodium channel function or perinexal structure directly affect cardiac conduction by altering the rate of cellular depolarization and ephaptic communication. Specifically, it is well established that sodium channel blockade or loss of function slows conduction.^7,8^ Additionally, previous work in animal models demonstrated that perinexal narrowing with dextran 2MDa, albumin, or hypercalcemia can accelerate conduction, whereas perinexal expansion with mannitol, hypocalcemia, or disruption of ID protein adhesion can slow conduction consistent with ephaptic mechanisms.^8–10^

Importantly, the auxiliary β-subunits of Na_V_1.5, the voltage-gated cardiac sodium channel, play important roles in channel trafficking, channel gating, Na_V_1.5 membrane localization and availability, and potentially ID adhesion.^9–11^ However, the role of β-subunit expression on peak sodium current (INa) density is less clear. For example, β1 co-expression with Na_V_1.5 can increase peak INa density,^12^ while full loss of β1 expression can also increase peak INa density.^13,14^ Additionally, homozygous null SCN1B mice display perinexus-specific ID de-adhesion relative to wild-type littermates.^9^ Taken together, β1 may facilitate ephaptic communication by maintaining nanoscale intercellular architecture and adhesion within the perinexus.

Clinically, mutations in SCN1B have been identified in patients with febrile seizures and epilepsy, Dravet syndrome, Brugada syndrome, and developmental and epileptic encephalopathy.^15–18^ Importantly, these disorders are associated with monoallelic or biallelic SCN1B mutations, not complete loss of β-subunits.^14^ Furthermore, SCN1B variants identified in patients often demonstrate incomplete penetrance,^19^ suggesting that partial loss of β1 subunit function may confer a latent or stress-dependent vulnerability rather than a constitutive phenotype. The majority of previous studies have primarily utilized a homozygous null Scn1b model, while more recent monoallelic and biallelic variants have also been studied.^20^ Regardless of the genetic background, no studies have reported conduction slowing, suggesting either a concealed phenotype or a negligible effect of Scn1b expression on cardiac conduction.^9,20,21^ On the other hand, peptide studies designed to competitively antagonize Scn1b adhesion have reported slowed conduction.^9^ Nevertheless, to our knowledge, no studies have appropriately recapitulated the phenotype of a clinically relevant conduction disease associated with Scn1b loss-of-function. Furthermore, the cardiac nano-structural and functional phenotype of a heterozygous-null model, which may more closely resemble the partial loss-of-function experienced by those with SCN1B mutations, has not been well-characterized.

The purpose of this study is to test the general hypothesis that Scn1b can be associated with a concealed conduction phenotype, which would be modulated by disruption to ephaptic coupling. In this study, we characterize the nano-structural and electrophysiologic phenotype of a heterozygous-null SCN1B model from mRNA to *in vivo* electrophysiology and hypothesize that a partial loss of SCN1B will confer increased sensitivity to modulation of perinexal structure and conduction via osmotic stress.

By focusing on a heterozygous null Scn1b model, this study addresses a gap between existing homozygous loss-of-function models and the genetic backgrounds observed in patients with SCN1B-associated disease. The combined nano-structural and functional analyses presented here demonstrate how a partial loss of β1 alters ID nanostructure and modulates cardiac conduction in response to osmotic stress. In doing so, this work implicates partial loss of β1-mediated adhesion in shaping ID nanostructure and conduction sensitivity, consistent with a mechanism for latent, context-dependent arrhythmia susceptibility.

## METHODS

All protocols were carried out in accordance with the principles of the Basel Declaration and conformed to the Guide for the Care and Use of Laboratory Animals published by the National Institutes of Health. The protocol was approved by the Institutional Animal Care and Use Committee of Virginia Polytechnic Institute and State University.

### Mouse Langendorff Perfusion

12-24-week-old 129-C57BL/6 Scn1b+/− mice of both sexes and age-matched, wild-type (WT) littermates, provided by Dr. Lori Isom, were euthanized via inhalant isoflurane and cervical dislocation. Hearts were excised and swiftly cannulated onto a Langendorff system as previously described.^8^ Hearts were retrogradely perfused with a crystalloid solution, bubbled with 100% oxygen, consisting of (in mM): 142.8 NaCl, 1.8 CaCl_2_, 6.0 NaOH, 4 KCl, 5.6 Dextrose, 1 MgCl_2_, 10 HEPES, and 1.2 NaH_2_PO_4_. pH was adjusted to 7.4 at 37°C using HCl (1N) or NaOH (1N). Hearts were immersed in a three-dimensional printed polylactic acid bath filled with crystalloid solution maintained at 37°C.^22^ Perfusion pressure was maintained (60-80mmHg) by variable flow (1.0-1.5mL/min) of perfusate.

### Optical Mapping

After a stabilization period of 10-15 minutes, hearts were injected through an injection port with a bolus of the voltage-sensitive dye, di-4 ANEPPS (60μM, Biotium, Inc.), followed by a 15-minute washout period in which the electromechanical uncoupler blebbistatin (10μM, Sigma-Aldrich) was used to reduce motion artifacts.

A 0.125mm unipolar silver-chloride electrode was placed on the anterior left ventricular epicardium and a ground electrode was placed in the bath. The heart was paced with a pulse width of 2ms and a basic cycle length of 150ms at 1.5X the diastolic threshold, as previously described.^23^ Hearts were paced for data collection but otherwise allowed to beat intrinsically.

The fluorophore was excited with a LED light source with fiber optic guide and filtered through an excitation filter (520/35nm), and emitted light passed through a dichroic mirror (565nm), and emission filter (610nm long-pass) before being detected by a MiCam Ultima L-type CMOS camera (SciMedia: 100×100 pixels, M=1.6X or 1.0X using a tandem lens configuration (back lens = 1.0X, forward lens = 1.6X or 1.0X, FOV = 9.9 x 9.9mm). Action potentials were recorded at a 1kHz sampling rate for ∼2s.

Hearts were treated with one of three interventions, which were perfused for 15 minutes following a baseline recording: time control with baseline solution (N= 8 WT, 4 Scn1b+/−), perinexal widening with mannitol (26.1g/L, N= 8 WT, 8 Scn1b+/−), or perinexal narrowing with dextran 2MDa (10g/L, N= 5 WT, 6 Scn1b+/−). After optical mapping experiments, hearts were subsequently utilized for tissue collection. Optical maps and all subsequent tissue were excluded from data analysis if insufficient dye loading occurred, marked by non-uniform staining in one or both ventricles.

### Tissue Collection

After perfusion for optical mapping experiments, atria were removed and left ventricle was separated out for tissue collection. A 1mm wide strip of left ventricular tissue from the anterior free wall was cut into ∼1mm^3^ pieces and fixed in a 4% paraformaldehyde, 2.5% glutaraldehyde, 1X phosphate-buffered saline (PBS) solution overnight at 4°C. The remaining left ventricular tissue was flash frozen in liquid nitrogen and stored at - 80°C.

### RNA purification and qRT-PCR

RNA was isolated from snap-frozen left ventricles. Tissues were lysed in Trizol (ThermoFisher #15596026) per manufacturer’s protocol and separated into phases using Phase lock gel tubes (Fisher #NC1093153). The top clear aqueous layer was further purified using the NucleoSpin RNA Kit (Macherey-Nagel #740955), and cDNA was generated using the SuperScript IV First-Strand Synthesis System (ThermoFisher #18091050), according to the manufacturer’s instructions. cDNA was used at a concentration of 20ng per qRT-PCR reaction with gene specific primers (Scn1b forward: 5’ – TCA CCT ACA ACC ACT CTG GCG AC- 3’; Scn1b reverse: 5’ – TGC CAT ATC TCT GTT GGC CTT GTC C-3’; GAPDH forward: 5’-AAT GGT GAA GGT CGG TGT G-3’, GAPDH reverse: 5’-GTG GAG TCA TAC TGG AAC ATG TAG-3’) and SYBR Select Master Mix for CFX (ThermoFisher #4472937). All samples were normalized to the threshold values of GAPDH as the housekeeping gene and data are represented relative to WT expression.

### Western Blots

Western blotting was performed using snap-frozen left ventricles. Briefly, the samples were homogenized in RIPA lysis buffer (containing 50mM Tris pH 7.4, 150mM NaCl, 1mM EDTA, 1% Triton X-100, 1% sodium deoxycholate, 0.1% sodium dodecyl sulfate, 200μM Na3VO34, 1mM NaF, and 5.6mM N-Ethylmaleimide) supplemented with Roche Protease Inhibitor Cocktail (Millipore Sigma #4693159001). Protein concentration was determined by a BioRad DC protein assay (#500-0112) and concentrations were normalized prior to analysis. Electrophoresis was performed to separate proteins which were then transferred to a PVDF membrane, blocked with 5% bovine serum albumin or 5% non-fat milk for 1 hour at room temperature and incubated overnight with a primary antibody against either the principal ventricular gap junction protein Cx43 phosphorylated at Ser368 (pCx43-S368, Cell Signaling Technology #3511S, validated in Solan et al.^24^, 1:1000 dilution), the voltage-gated sodium channel β1 subunit (D4Z2N, Cell Signaling Technology #13950, 1:1000 dilution, validated in Williams, Z. et al,^25^) or against the voltage-gated sodium channel protein Na_V_1.5 (Cell Signaling Technology #14421s, validated in Blair, G. et al.,^23^ 1:1000 dilution), at 4°C. The membranes were then washed and incubated with secondary antibody (Abcam #ab6721, 1:10,000 dilution) at room temperature for 1 hour. After washing, bound antibody was detected using West Pico Plus chemiluminescent substrate (ThermoFisher #34579) and imaged using the Licor Odyssey Fc system. Membranes were stripped with ReBlot Plus (Millipore Sigma #2504) according to manufacturer’s instructions, blocked in Intercept Blocking Buffer (Licor #927-60001) at room temperature for 1 hour and incubated with primary antibodies against total Cx43 (Millipore Sigma #C6219, validated in George, S. et al.,^26^ 1:5000 dilution) and/or Gapdh (Santa Cruz #sc-47724, 1:5000 dilution).

Membranes were then washed and incubated with secondary antibodies for 1 hour (Licor #925-32211, 1:10,000 dilution; and Licor #925-68070, 1:10,000 dilution) and washed again. Membranes were again imaged using the Licor Odyssey Fc system to determine protein expression. Total Na_V_1.5, β1-subunit and Cx43 protein expression were normalized to Gapdh and pCx43-S368 expression was normalized to the total Cx43 expression.

### Myocyte Isolation

Ventricular myocytes from WT (N=4) and Scn1b+/− (N=4) mice were isolated using methods adapted from a previous study.^27^ The primary perfusion buffer used for cell isolation contained, in mM: 130 NaCl, 5 KCl, 0.5 NaH_2_PO_4_, 10 HEPES, 1 MgCl_2_, 10 Taurine, 10 Glucose. pH was adjusted to 7.80 at room temperature using NaOH. The perfusion buffer was supplemented with 15 μM blebbistatin (Cat#Β1387, APExBIO) at each step of the myocyte isolation procedure. Additionally, 1.25mg/mL collagenase type 2 (285μL/mg dw, Cat#LS004176, Worthington Biochemical Corporation), or fetal bovine serum (Gibco Cat#A5670401, Thermo) to block the effect of the enzyme, were added during the tissue digestion step, as detailed below. Mice were euthanized by isoflurane (Fluriso,VetOne) overdose and cervical dislocation, followed by immediate midsternal thoracotomy and removal of the pericardium to expose the heart. After rapidly cutting the inferior vena cava, excess blood was cleared using a 27-gauge needle syringe to inject 4-7mL of perfusion buffer into the right ventricle at approximately 2mL/min. The heart was then removed and the aorta was clamped, followed by injection of an additional 7-9mL of perfusion buffer into the left ventricle via the apex at approximately 2mL/min. To digest the heart, 25-35mL of prewarmed (37°C) collagenase type 2-containing perfusate was injected via the same apical orifice at approximately 2mL/min. Once the tissue was digested, the ventricular tissue was separated out and placed into 4mL of perfusion buffer. Tissue samples were then gently torn into smaller pieces using forceps. Perfusion buffer containing fetal bovine serum to initiate block of the digestive enzyme was added until the total volume equaled 10mL. The solution containing the tissue was then triturated for 2-3 minutes using a transfer pipette, after which remaining pieces of intact tissue were discarded, leaving only myocytes in solution. The enzyme-containing solution was gradually exchanged for fetal bovine serum-containing solution using a centrifuge (Sorvall Legend X1R, Thermo Fisher Scientific) at 500G to separate the myocytes from the enzyme-containing solution over 2-3 iterations. Once digestion was fully blocked, extracellular Ca^2+^ was elevated in a stepwise manner to 300μM, 600μM, and finally 1mM, with 6-9 minutes between steps. The solution with myocytes was then placed over laminin-covered 8mm diameter cover slips for use on the microscope stage of the patch clamp setup.

### Patch Clamp Recording of Sodium Current

INa was recorded using a voltage-clamp setup which included an Axopatch 200B amplifier (Molecular Devices, LLC), an Axon Digidata 1550B digitizer (Molecular Devices, LLC), and an inverted microscope (Olympus IX83, Evident Scientific). A glass microelectrode puller (Narishige PC-10) was used to fabricate borosilicate glass (1.5mm OD/1.12mm ID, World Precision Instruments) microelectrodes (tip resistance 2-4 MΩ). Using micromanipulators (Scientifica), the microelectrodes were directed to the lateral membrane of the myocyte to form a ‘gigaseal’ and implement a whole-cell voltage clamp configuration. Leak current was subsequently subtracted and series resistance was compensated to greater or equal to 70%. Patched cells were subject to a voltage clamp protocol (−100mV holding potential with 300ms test pulses progressing from - 80mV to +30mV in 5mV increments), during which triggered currents were low-pass filtered (2kHz) and recorded at a 20kHz sampling rate using the voltage clamp amplifier/digitizer system and pClamp software (Molecular Devices, LLC). INa recordings were carried out at room temperature in a bathing ‘external’ solution containing (in mM): 25 NaCl, 120 N-methyl-D-glucamine, 10 HEPES, 1.8 MgCl_2_, 1.8 CaCl_2_, and 10 Glucose. pH was adjusted to 7.3 with HCl. The external solution was supplemented with 100µM verapamil hydrochloride (Cat#V4629, Sigma Aldrich) to block calcium currents. The composition of the microelectrode ‘internal’ solution contained (in mM): 120 CsMeSO_3_, 20 Tetraethylammonium chloride, 2 MgCl_2_, 10 HEPES, 10 EGTA, and 4 Mg-ATP. The pH was adjusted to 7.3 using CsOH. The magnitude of the peak current at each test potential was determined off-line using pClamp software. Current density values (pA/pF) were computed by normalizing the measured currents (pA) to the cellular membrane capacitance (pF) determined for each myocyte during voltage clamp.

### Transmission Electron Microscopy

Fixed left ventricular samples were post-fixed with 2% Osmium Tetroxide for 1 hour on ice, followed by en bloc staining with 1% Uranyl Acetate in 25% ethanol for 1 hour.

Samples were then dehydrated through an ethanol series (50%, 70%, 90%, and 100%). Samples were infused with Embed 812 resin (EMS #14900) (propylene oxide/Embed 812, 50:50) overnight, followed by 4 changes of 100% resin. Tissue samples were then embedded in BEEM capsules and Embed 812 resin, polymerized for 48 hours at 65°C. 70nm, gold, ultra-thin sections were cut using a Leica EM UC7 ultramicrotome and electron micrographs were produced with an FEI Tecnai G2 Spirit Biotwin transmission electron microscope at 120 kV. Images were taken at 18,500X magnification.

### In Vivo Assessment

Mice were anesthetized using 5% inhalant isoflurane (MWI, Fluriso) and 3L/min oxygen in an airtight induction chamber, then maintained at 3% isoflurane and 4L/min oxygen on a nose cone. After lack of reflexes was observed, needle electrodes were inserted into the forelegs and left hindleg of the animal as previously described^28^ and electrocardiogram (ECG) was recorded for 3 minutes at 1kHz sampling rate. Mice were then retro-orbitally injected with 150μL of either 0.9% NaCl (MWI Animal Health; N= 5 WT, 5 Scn1b+/−) or 26.1g/L mannitol dissolved in 0.9% NaCl (N= 5 WT, 5 Scn1b+/−).

After 10 minutes of recovery on a heating pad, mice were re-anesthetized, needle electrodes were reinserted, and ECG was recorded for 3 minutes for paired baseline to intervention measurements.

### Data and Statistical Analysis

Optical signals were binned 2×2, resulting in an interpixel resolution of 0.12mm for 1.6X magnification and 0.198mm for 1.0X magnification. Action potential activation times were assigned as previously described.^29^ Conduction velocity was estimated in the transverse and longitudinal directions using a semi-automated custom MATLAB program.^30^ The following parameters were used for assigning activation time: length of boxcar filter: 3, number of consecutive sample points defining a peak: 3, blockout interval after detection of peak: 60ms, peak amplitude and direction multiplier factor: - 1.1. Vectors were assigned using the Bayly algorithm^29^ with the following parameters: distance between mapping sites – X: 0.12mm or 0.198mm, distance between mapping sites – Y: 0.12mm or 0.198mm, distance over which vectors are calculated: 4 pixels, time over which vectors are calculated: 200ms, minimum number of sites to include: 12, maximum allowable error for each vector: 0.8, minimum allowable vector magnitude: 0.01 m/s, maximum allowable vector magnitude: 1 m/s. Masking parameters included vectors up to dilation 14 (1.68mm at 1.6X or 2.772mm at 1.0X) away from the pacing site and within an angle of 16 degrees.

A one-sample t-test compared Scn1b+/− mRNA expression to a value of 1 for WT. Western blot images were analyzed using Li-Cor Image Studio and compared using unpaired t-tests. Transmission electron micrographs were analyzed as previously described^31^ to quantify perinexal width. Perinexal width was compared using ordinary two-way ANOVA. Conduction velocity was compared using Repeated Measures (RM) two-way ANOVA, and the change in conduction velocity (ΔCV) was compared using unpaired t-tests. ECG parameters were analyzed using the built-in ECG Analysis in LabChart and the entire 3-minute recording was averaged to return one QRS duration measurement per mouse per condition. QRS was compared using RM two-way ANOVA. All statistics completed in GraphPad Prism v9.0 or later. P < 0.05 is considered statistically significant. Data are presented as mean and standard deviation, unless otherwise noted.

## RESULTS

### Heterozygous loss of Scn1b reduces β1 expression without altering canonical conduction proteins

As confirmed with RT-PCR, there was approximately a 50% reduction in Scn1b mRNA expression between the wild-type (WT) and Scn1b+/− mice (Figure 1A). Western blot was performed (Figure 1B) to assess β1 protein expression (Figure 1C), demonstrating a significant reduction in β1 protein levels between WT and Scn1b+/− mice. Canonical conduction protein expression was also verified, to ensure that Na_V_1.5, connexin-43, and phosphorylated connexin-43 were not affected by the reduction in β-subunits.

**Figure 1:**
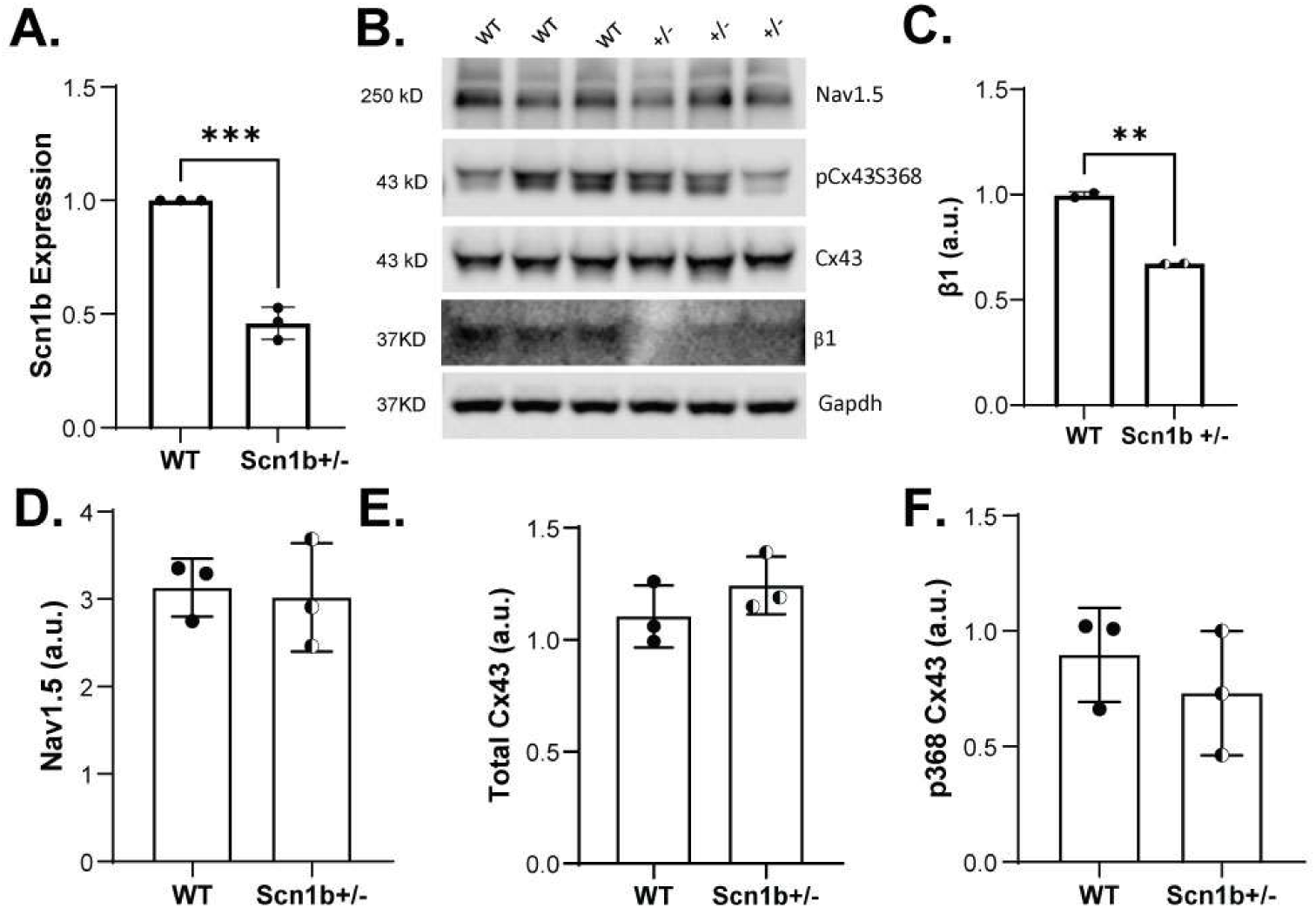
Scn1b+/− mice have reduced Scn1b mRNA and β1 protein, with no change in canonical conduction protein expression. **A.** Quantification from RT-PCR demonstrating a significant, ∼50% reduction in Scn1b mRNA between WT and Scn1b+/−mice. *, P < 0.05, one-sample t-test. **B.** Representative Western Blot image from left ventricular tissue. **C.** Quantification of β1 expression from two technical replicate Western Blots (n=2) with three biological replicates (N=3) each, demonstrating a significant reduction in β1 protein in Scn1b+/− hearts relative to WT, normalized to GAPDH. **D.** Quantification of Na_V_1.5 expression from Western Blot with three biological replicates (N=3) showing no difference between WT and Scn1b+/−, normalized to GAPDH. **E.** Quantification of Total Cx43 expression from Western Blot (N=3) showing no difference between WT and Scn1b+/−, normalized to GAPDH. **F.** Quantification of serine-368-phosphorylated Cx43 from Western Blot (N=3) showing no difference between WT and Scn1b+/−, relative to total Cx43. *, P < 0.05, unpaired t-test.

There was no significant difference in Na_V_1.5 expression normalized to GAPDH (Figure 1D), total connexin-43 expression normalized to GAPDH (Figure 1E), or the ratio of S368-phosphorylated connexin-43 normalized to total connexin-43 (Figure 1F) in Scn1b+/− relative to WT mice.

### Peak sodium current is preserved in Scn1b+/− ventricular myocytes

Representative sodium current (I_Na_) traces from WT and Scn1b+/− ventricular myocytes are shown in Figures 2A and B respectively, as measured by whole-cell patch clamping. The I-V curves generated by averaging all recordings of each group (Figure 2C) and the groupwise average peak current (Figure 2D) demonstrate no significant I_Na_ difference between WT (−66.62 pA/pF) and Scn1b+/− (−60.35 pA/pF) ventricular cardiomyocytes.

**Figure 2:**
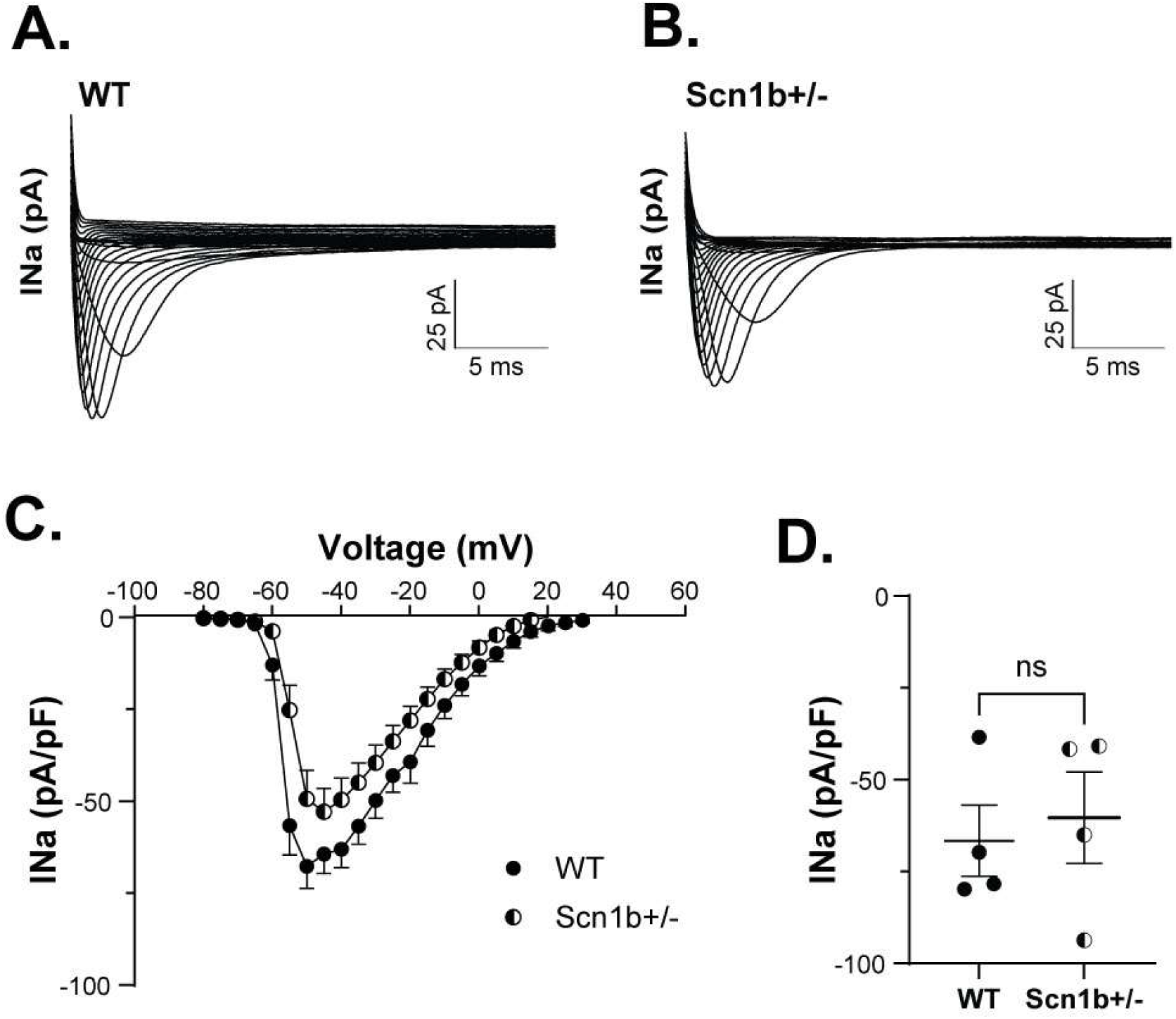
Ventricular I_Na_ is preserved between WT and Scn1b+/− cardiomyocytes. **A.** Representative I_Na_ traces from a WT cardiomyocyte. **B.** Representative I_Na_ traces from a Scn1b+/− cardiomyocyte. **C.** Average I-V curves from WT (N=3 hearts, n=21 cells) and Scn1b+/− (N=4 hearts, n=18 cells) cardiomyocytes. Data is presented as mean and standard error of the mean. **D.** Groupwise quantification of I_Na_ peak current, i.e. the average value of the mean peak I_Na_ in each heart, demonstrating no difference between WT and Scn1b+/− cardiomyocytes. Unpaired t-test.

### Partial loss of Scn1b alters perinexal structure and its response to osmotic challenge

Perinexal structure was quantified by measuring perinexal width (W_P_). Transmission Electron Microscopy (TEM) was used to measure W_P_ under each of the three conditions (baseline, mannitol, and dextran) in both genotypes (Figure 3A). At baseline, Scn1b+/−hearts (25.3 nm) displayed a larger mean W_P_ than WT hearts (20.9 nm). In WT hearts, mannitol (29.2 nm) significantly increased W_P_ relative to baseline, but dextran (20.9 nm) did not cause a detectable change in W_P_ (Figure 3B). In Scn1b+/− hearts, mannitol (29.4 nm) significantly increased W_P_, while dextran 2MDa (19.2 nm) significantly reduced W_P_ (Figure 3B).

**Figure 3:**
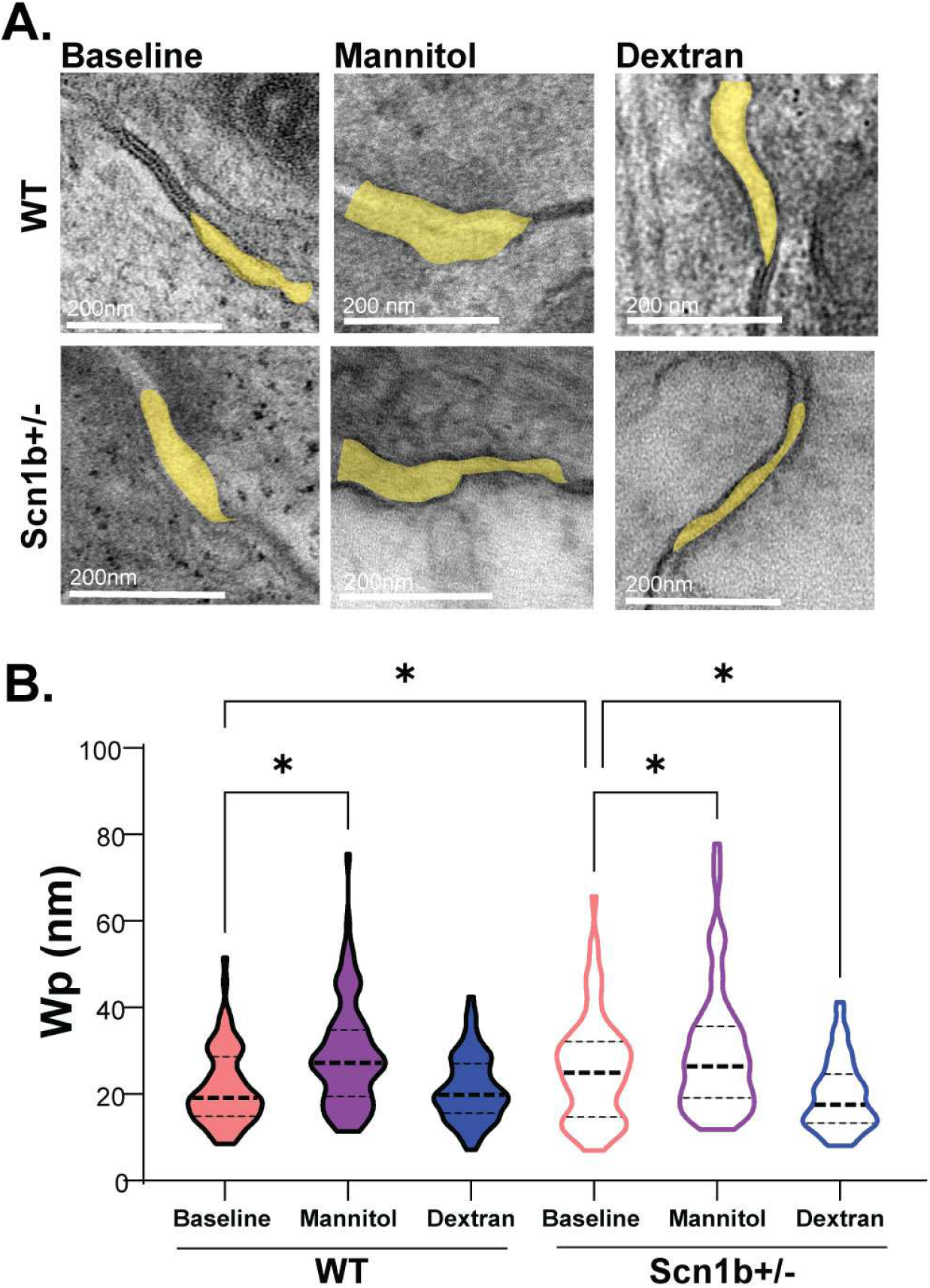
W_P_ is wider in Scn1b+/− hearts, and more sensitive to osmotic stress. **A.** Representative perinexal tracing (yellow) from WT (*top*) and Scn1b+/− (*bottom*) hearts under the three interventions: baseline *(left*), mannitol *(center*), and dextran *(right*). **B.** Quantification of perinexal width (W_P_) from N=4 hearts per intervention, with n=15-20 perinexal images per heart. Scn1b+/− hearts have wider perinexi at baseline relative to WT. Mannitol significantly increases W_P_ in both genotypes. Dextran 2MDa selectively narrows perinexi in Scn1b+/− hearts. *, P < 0.05, Ordinary two-way ANOVA.

### Scn1b+/− hearts have preserved conduction at baseline, but increased sensitivity to osmotic stress ex vivo

In pooled baseline data, there was no significant difference in conduction velocity as measured *ex vivo* with optical mapping between WT (40.8 cm/s) and Scn1b+/− (42.4 cm/s) hearts (P=0.5037, Supplemental Figure 1A). When baseline recordings were compared with paired fifteen-minute continued perfusion time controls, there were no significant changes to conduction velocity in either WT (baseline: 38.6 cm/s, time control: 40.6 cm/s) or Scn1b+/− (baseline: 39.0 cm/s, time control: 39.2 cm/s) hearts (Figure 4A-C). Both WT (baseline: 42.4 cm/s, mannitol: 37.4 cm/s) and Scn1b+/−(baseline: 48.5 cm/s, mannitol: 38.8 cm/s) hearts displayed significant conduction slowing in response to 15 minutes of perfusion with 26.1g/L mannitol (Figure 4D-E); furthermore, mannitol had a significantly greater effect on Scn1b+/− hearts than WT hearts (−9.7 cm/s compared to –5.0 cm/s, Figure 4F). Perfusion with 10g/L dextran 2MDa resulted in no change in conduction velocity in the WT (baseline: 38.3 cm/s, dextran: 37.1 cm/s) hearts, whereas it led to a mild but significant increase in conduction velocity in Scn1b+/− (baseline: 34.7 cm/s, dextran 36.7 cm/s) hearts (Figure 4G-I).

**Figure 4:**
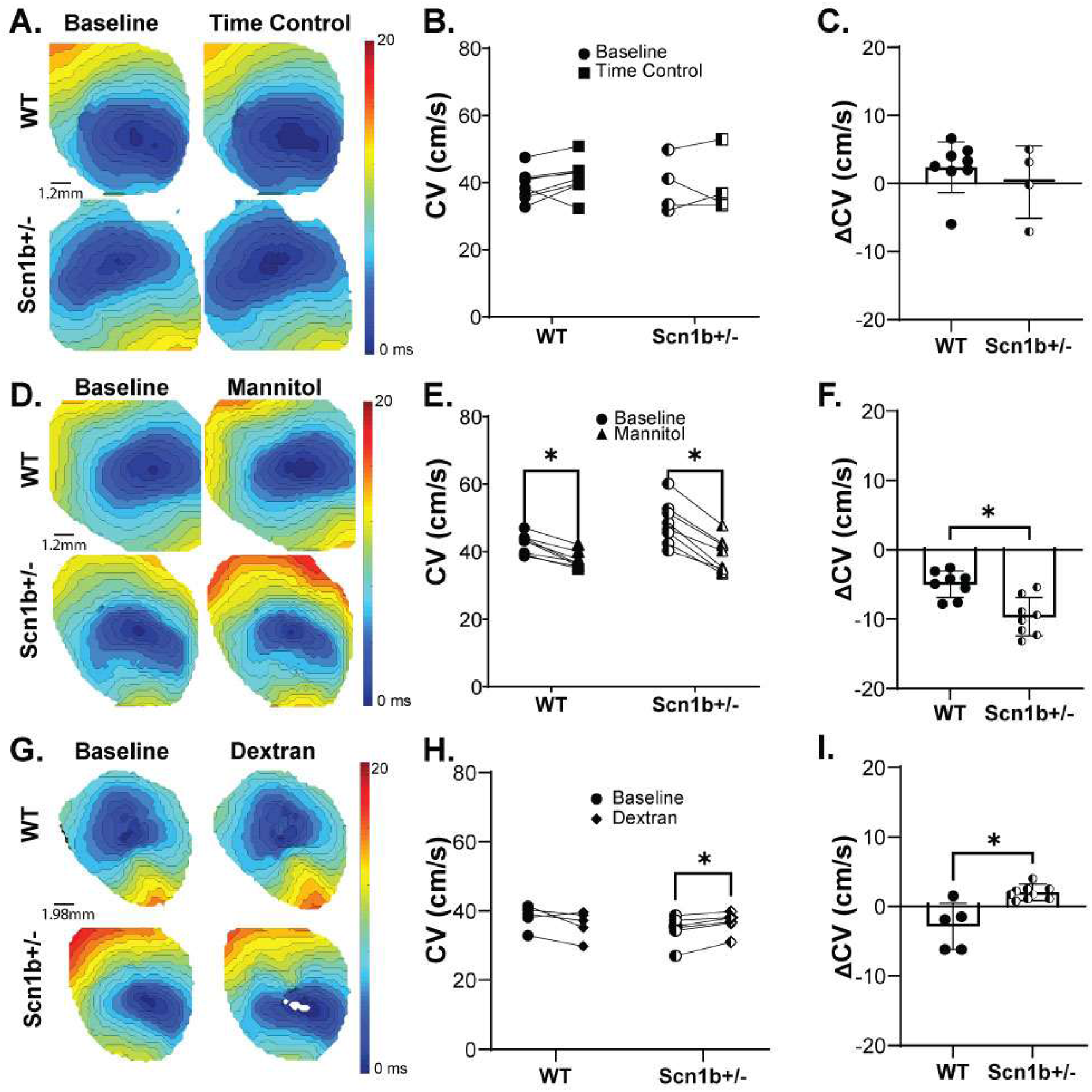
Scn1b+/− hearts have preserved conduction at baseline but are more sensitive to osmotic stress. **A.** Representative isochrone activation maps of WT (*top,* N=8) and Scn1b+/− (*bottom,* N=4) hearts at baseline (*left*) and after 15-minutes continued perfusion for time control (*right*). **B.** Quantification of CV in the transverse direction for WT and Scn1b+/− hearts at baseline and time control demonstrating no significant differences. **C.** Delta CV (time control – baseline) for WT and Scn1b+/− hearts. **D.** Representative isochrone activation maps of WT (N=8) and Scn1b+/− (N=8) hearts at baseline and after 15-minutes perfusion of 26.1g/L mannitol. **E.** Quantification of CV in the transverse direction for WT and Scn1b+/− hearts at baseline and mannitol demonstrating mannitol significantly slows CV in both genotypes. **F.** Delta CV (mannitol – baseline) for WT and Scn1b+/− hearts demonstrating mannitol slows CV to a significantly greater degree in Scn1b+/− hearts. **G.** Representative isochrone activation maps of WT (N=5) and Scn1b+/− (N=6) hearts at baseline and after 15-minutes perfusion of 10g/L dextran 2MDa. **H.** Quantification of CV in the transverse direction for WT and Scn1b+/− hearts at baseline and dextran demonstrating dextran speeds CV only in Scn1b+/− hearts. **I.** Delta CV (dextran – baseline) demonstrating a significant difference in the effect of dextran between genotypes. *, P < 0.05, RM two-way ANOVA; unpaired t-test.

### Scn1b+/− hearts are sensitive to osmotic stress in vivo

A representative trace of sinus rhythm recorded from a Scn1b+/− mouse at baseline is presented in Figure 5A. Interestingly, one Scn1b+/− mouse developed an arrhythmia after injection with mannitol, of which a representative trace is provided in Figure 5B. Representative average QRS traces from each of the two genotypes under three conditions are presented in Figure 5C, demonstrating that WT and Scn1b+/− mice had similar QRS durations at baseline. WT mice displayed no significant changes to baseline QRS duration after either normal saline (baseline: 11.0 ms, saline: 11.0 ms) or mannitol (baseline: 11.0 ms, mannitol: 11.5 ms) injection (Figure 5D, *left*). Conversely, Scn1b+/− mice displayed no significant changes to baseline QRS duration after normal saline (baseline: 10.8 ms, saline: 10.8 ms) injection, but significant prolongation of QRS after mannitol (baseline: 10.5 ms, mannitol: 13.2 ms) injection (Figure 5D, *right*).

**Figure 5:**
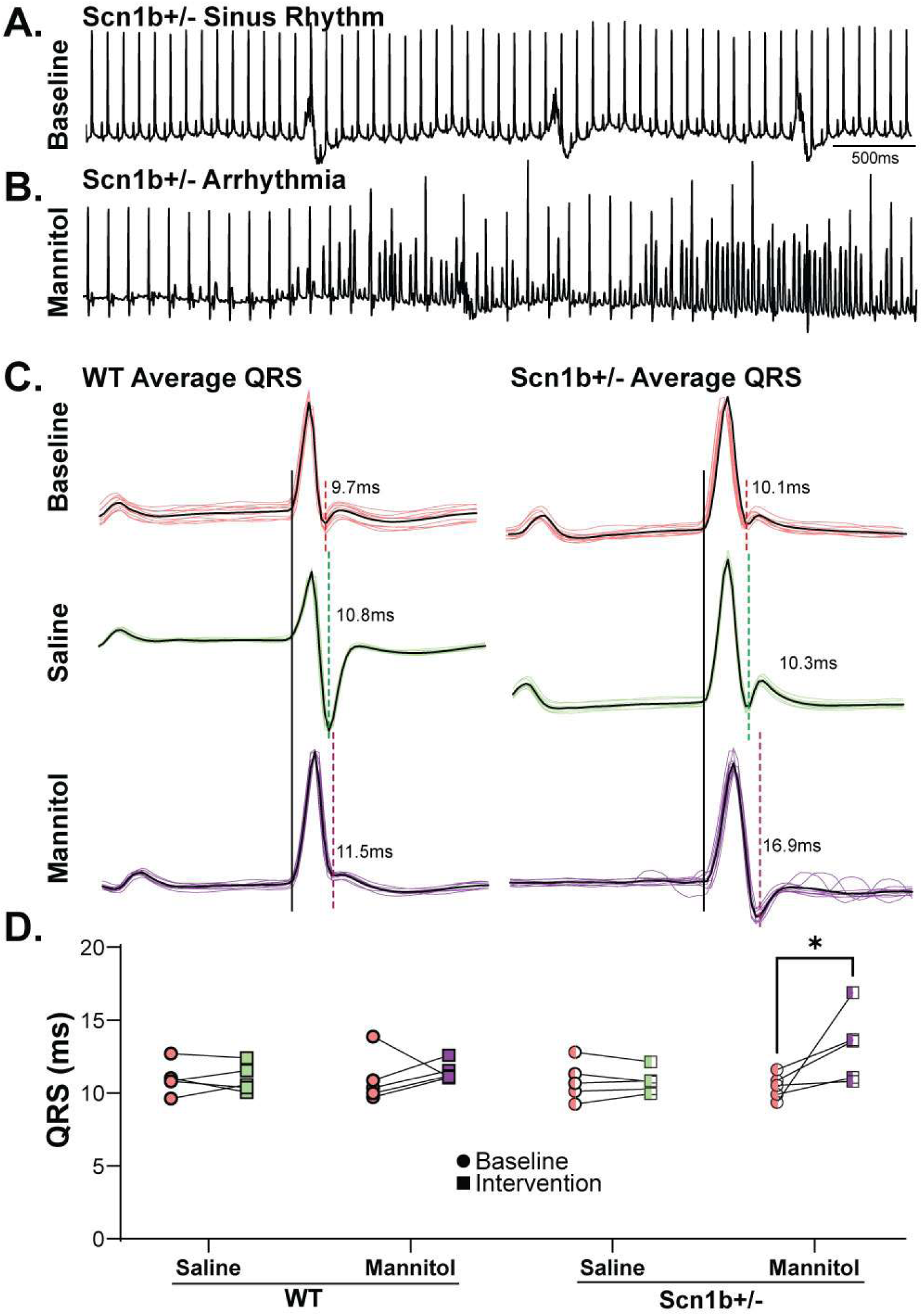
Concealed phenotype and conduction sensitivity to osmotic challenge are preserved *in vivo*. **A.** Representative trace of 5s ECG recording of sinus rhythm in Scn1b+/− mouse. **B.** Exemplar trace of 5s ECG recording of arrhythmia in Scn1b+/− mouse after mannitol injection. **C.** Average QRS traces (black) from n=10 traces (color) for WT (*left*) and Scn1b+/− (*right*) mice after each intervention: baseline (*top*), 0.9% saline injection (*center*), and 26.1g/L mannitol injection (*bottom*). **D.** Quantification of average QRS durations from entire 3-minute recording for WT before and after saline (N=5) or mannitol (N=5) injection and for Scn1b+/− before and after saline (N=5) or mannitol (N=5) injection demonstrating significant prolongation of QRS duration in Scn1b+/− mice after mannitol injection. *, P < 0.05, paired t-test.

## DISCUSSION

In this study, we define the cardiac nano-structural and electrophysiologic phenotype associated with heterozygous loss of the sodium channel β1 subunit encoded by Scn1b. Using a multiscale approach spanning molecular expression, cellular electrophysiology, intercalated disc ultrastructure, *ex vivo* conduction, and *in vivo* electrocardiography, we demonstrate that β1 haploinsufficiency produces a latent conduction vulnerability that emerges secondary to intercalated disc dehiscence induced by osmotic stress.

At the molecular and cellular levels, heterozygous deletion of Scn1b resulted in an approximately 50% reduction in β1 mRNA and protein, without detectable changes in expression of Na_V_1.5, connexin-43 (Cx43), or Cx43 phosphorylation at serine 368. This is consistent with previous work, which established that there is no difference in Na_V_1.5 mRNA between wild-type (WT) and homozygous null Scn1b mutants,^32^ but, to our knowledge, this is the first demonstration that heterozygous null Scn1b does not measurably alter Na_V_1.5 or Cx43 protein expression.

Furthermore, peak sodium current (INa) was not significantly different between WT and Scn1b+/− in isolated ventricular myocytes, demonstrating that a 50% loss of β1 does not alter functional Na_V_1.5 expression in this model. However, the relationship between β1 and Na_V_1.5 functional expression in previous studies is conflicting. Specifically, some groups report over-expressing β1 increases Na_V_1.5 functional expression,^13^ while others report the opposite; knocking out Scn1b increases peak INa.^14^ Furthermore, two additional studies demonstrated that a monoallelic β1 mutant decreased peak INa,^20,33^ and one of those revealed the biallelic mutant did not change INa.^33^ Thus, Scn1b complexly modulates functional Na_V_1.5 expression, which could be attributable to experimental differences. For example, some groups used heterologous expression systems while others used isolated ventricular myocytes from transgenic mice.

Furthermore, knocking out the protein is different from genetic mutations, and genetic mutations can confer very specific phenotypic differences based on the location of the mutation within the protein. Since β1 is implicated in many different functions associated with Na_V_1.5, including trafficking, surface expression, post-translational modification, and gating, there is the opportunity for highly variable effects based on the specific mutation studied. Regardless, a partial loss of Scn1b in a transgenic mouse model does not alter the protein expression of canonical determinants of cardiac conduction or functional Na_V_1.5 expression. Importantly, changes to INa in isolated ventricular myocytes do not always translate to a tissue or global conduction phenotype.

At the next larger scale of cellular networks, we previously demonstrated that a conduction phenotype can arise from mechanisms predicated on intercellular architecture.^34^ The suggested mechanism is that reduced intercalated disc adhesion, leading to widening in the gap junction adjacent perinexus, which densely expresses Na_V_1.5, can reduce a specialized form of cardiac conduction called ephaptic coupling.^9,35,36^

Transmission electron microscopy in the present study revealed that baseline perinexal width (W_P_) is wider in Scn1b+/− hearts relative to WT hearts, consistent with previous findings that perinexi are wider in Scn1b-/- murine hearts,^9^ and claims that β1 is important for intercalated disc adhesion.^25^ While a direct comparison of W_P_ in Scn1b+/−hearts (∼25nm) appears to be close to half of W_P_ reported in a previous study with Scn1b-/- mouse hearts (∼40nm), direct comparisons should be interpreted cautiously as we demonstrated manual segmentation can produce analyst dependent absolute values, even if different analysts can report similar and significant W_P_ changes with intervention or disease.^37^

Despite modest but significant W_P_ differences at baseline, conduction velocity was not different between WT and Scn1b+/− hearts. The lack of a baseline conduction phenotype in Scn1b+/− hearts is consistent with previous studies in the presumably more severe Scn1b-/- models that also did not report significant conduction slowing.^14,38^

Importantly, clear genotype-dependent conduction differences emerged when intercalated disc structure was challenged with osmotic stress. Perfusion with mannitol slowed conduction in both wild-type and Scn1b+/− hearts, consistent with previous mouse heart optical mapping experiments,^23^ but the magnitude of slowing was significantly greater in Scn1b+/− hearts. This demonstrates that the conduction phenotype in Scn1b+/− hearts can be concealed under baseline conditions and unmasked with the osmotic agent mannitol.

Importantly, computational models of ephaptic coupling predict a complex bi-phasic association between cardiac conduction and W_P_.^39^ Specifically, conduction velocity can be invariant over some nominal range of ephaptic and gap junction coupling when self-activation and self-attenuation are balanced. Widening clefts beyond that transition zone, even modestly, can dramatically slow conduction by further reducing transactivation, and gap junction conductance.^55^ Thus, β1 haploinsufficiency may insufficiently widen the perinexus at baseline conditions, but a further increase in W_P_ with mannitol can dramatically slow conduction, indicating a latent, context-dependent vulnerability that is unmasked under stress.

To further explore this effect, hearts were treated with dextran 2MDa, which has been shown in guinea pigs to narrow the perinexus.^34^ Interestingly, dextran treatment produced no detectable change in W_P_ in WT hearts. Although this contrasts with prior findings, it may reflect a species-specific difference. Under baseline conditions, mouse intercalated discs may already be near their minimal spacing due to extracellular allosteric protein interactions. This interpretation suggests that WT mouse hearts may operate near a structural limit in which further narrowing does not enhance conduction.

In contrast, Scn1b+/− hearts exhibited significant perinexal narrowing following dextran treatment, accompanied by an increase in conduction velocity. Furthermore, treatment with mannitol increased W_P_ in both genotypes and was associated with a significant reduction in conduction velocity, with a more pronounced effect in Scn1b+/− hearts.

Together, these findings suggest the existence of a lower bound for W_P_, likely set by structural constraints within the intercalated disc. They also indicate that Scn1b+/−hearts display baseline intercalated disc dehiscence with a latent conduction phenotype that can be revealed under osmotic stress, supporting the idea that haploinsufficiency of Scn1b sensitizes cardiac conduction to perturbations in extracellular spacing.

The physiological relevance of this concealed phenotype is further supported by *in vivo* recordings. While baseline QRS duration was similar between genotypes, mannitol selectively prolonged QRS in Scn1b+/− mice. In one animal, mannitol triggered an arrhythmia during the recording period. Together with *ex vivo* data, these findings demonstrate that β1 haploinsufficiency produces a functional phenotype that is largely concealed under nominal conditions but becomes unmasked under osmotic perturbation.

Given that mannitol is used clinically, this study provides evidence that osmotic challenge can be a novel adjuvant or potential primary test for diagnosing concealed Brugada Syndrome. Importantly, the current gold standard for diagnosing Brugada Syndrome is flecainide challenge. However, this test has low sensitivity and a false-negative rate up to 32%.^40^ Given our findings, we propose further research into the safety and efficacy of osmotic challenge as a novel diagnostic test to unmask the Brugada phenotype. Additionally, these data suggest that patients with Brugada Syndrome may benefit from anti-inflammatory therapy to decrease arrhythmia vulnerability, as inflammation in the heart, particularly myocarditis, can lead to interstitial edema, similar to the effects of mannitol.^41^

Taken together, our results provide a coherent mechanistic framework: β1 subunits contribute to intercalated disc structural integrity, and partial loss of β1 sensitizes conduction to interventions and disease states that reduce ephaptic coupling. Consistent with previous studies, which also did not observe an abnormal conduction phenotype at baseline, Scn1b down-regulation or mutations associated with loss of functional adhesion in the intercalated disc are associated with invariant conduction relative to WT at baseline, but osmotic stress reveals a latent susceptibility. These observations have implications for understanding SCN1B-associated disease, where partial loss-of-function variants may exhibit incomplete penetrance: baseline conduction appears normal, but environmental or physiological stressors may precipitate conduction slowing or arrhythmia.

## LIMITATIONS

This study has several important limitations. First, while mice of both sexes were included, the study was not sufficiently powered to detect sex-specific differences in conduction or perinexal structure. Second, the age range of mice used (12-24 weeks) may not capture potential age-related changes in perinexal width or conduction; prior data in aged guinea pigs suggest age-related perinexal narrowing,^42^ though the animals in those studies were substantially older than those used here. Third, although we observed significant effects of osmotic challenge on perinexal structure and conduction, species- and strain-specific differences may limit direct extrapolation to humans, particularly regarding responses to dextran or mannitol. Finally, while one Scn1b+/−mouse developed an arrhythmia *in vivo*, the study was not sufficiently powered to make definitive claims regarding arrhythmia incidence or risk. Additionally, while we quantified total protein expression of canonical conduction determinants (Na_V_1.5, Cx43, and phosphorylated Cx43), we did not assess their precise subcellular localization. Prior studies indicate that β1 mutations can alter Na_V_,^43^ and it remains possible that subtle redistribution occurs without changes in total protein or INa.

